# cspray: Distributed Single Cell Transcriptome Analysis

**DOI:** 10.64898/2026.02.06.704110

**Authors:** Peter G. Hawkins, Eli M. Swanson, Megan Feichtel

## Abstract

The size of individual single cell samples continues to grow with advancing technologies, as do the number of samples included in individual experiments and across organizations. This presents challenges for processing this data at scale, both in terms of computational throughput and the required size of the machines that must process this data. We present a single cell RNA processing method that is fully distributed, capable of processing arbitrarily large files, and numbers of files, without requiring per-file based compute sizing. Our method, cspray, includes data ingestion, preprocessing, highly variable gene annotation, PCA, and clustering. We also show that this processing at scale permits LLM based reference-free cluster annotation on low resolution clusters, which demonstrates these techniques can be used to build single cell data discovery platforms at scale.

## INTRODUCTION

Droplet based single cell RNA processing began with samples of tens of thousands of cells [1, 2]. This has further been steadily increased with other technologies to maximize throughput and introduce multiplexing, see e.g Baysoy et al. [3]. Technologies such as PIPseq [4] are pushing these limits further still to above a million cells, and Elz et al. report commercial kits at up to 5 million cells today in their recent review of commercial single cell RNA sequencing technologies [5].

Tools such as Seurat [6, 7] and Scanpy [8] are invaluable for analyzing these data. However, with increasing file sizes, the pressure on the required RAM of the machines used for this processing rapidly grows. Further, the absolute number of cells processed by individual teams, organizations, and partnerships, continues to grow. This in itself requires the ability to process many samples on a regular basis. Therefore, efficient throughput of cells through any processing pipeline is increasingly important. When many samples are being processed at once, an additional challenge emerges: individual sizes of the files can vary, creating a difference in the compute size that must be taken into account if one does not want to over-provision the compute.

Several strategies for addressing these challenges are emerging. The most popular two approaches being: using GPUs for more efficient throughput, and using distributed compute. We comment further on these in the related work section. In this work we focus on the distributed compute approach, in particular in order to read arbitrarily large files and numbers of files at once, and process them, using distribution end to end.

When designing the cspray system for scalable, single cell transcriptomic analysis, we considered both technical features and some common challenges faced by scientists analyzing such datasets, so cspray might address them. Among the latter grouping, we identify two key challenges aside from the technical scalability challenges:

1. Scientists need to be able to identify samples outside of the current experiment for large scale integrations, and to do so, they need easy access to simple QC statistics, clustering, and cell type labeling to best select those
2. Scientists need to occasionally access the raw files as needed for deep ad-hoc analysis, and they should ideally be able to pick up results in intermediate stages, i.e. with basic QC already performed, or after PCA is performed, easily

On the first point, one may consider foundation models for single cells (see e.g. [9–12]) to be a perfect fit for cell type labeling across many samples. Indeed, running large scale inference of foundation models is inherently scalable. These models still typically require one to prepare the input data, which can involve out of memory (OOM) issues for large datasets. That is to say, we believe a scalable read and processing framework can form the basis of a large scale data preparation prior to inference with such models.

In this work, though, we focus instead on collecting high level cell annotations (e.g. T cell, B cell), rather than using single cell foundation models, as we believe these are already very useful for building large scale annotated datasets for data discovery. They are also efficient to compute at the cluster level [13, 14], and may be less susceptible to changes in sample quality and processing parameter options.

We thus sought to build a solution addressing both the technical and scientific challenges above, and we specifically focus on:

a. Distributed processing of data from raw files all the way to clustered data that scales on even small machines
b. Data written into efficient Delta™ format tables at many stages, including the raw data for efficient later access and operations
c. Focus on low resolution clustering that scales and pairs with reference-free cell type labeling at the cluster level

## RELATED WORK

In addition to the most popular scRNA analysis tools (e.g. Seurat [6, 7] and Scanpy [8]) used today that typically run on single CPU machines, there is growing interest in accelerating these workflows using GPUs and handling larger data sets using distributed compute.

The rapids-singlecell [15] library, part of NVIDIA^™^’s set of RAPIDS packages, was recently made a formal member of the scverse [16] set of packages and provides a much faster processing than the CPU based default scanpy [8] implementation. However, it is worth noting that one is still limited by the compute size and can experience OOM if one’s GPU is not big enough for any given sample. Similarly to scanpy, one may wish to also change compute size manually for different file sizes to control cost, requiring additional orchestration and overhead.

For the distributed approach, one hopes to split data among many workers to prevent OOM occurring on any individual machine while also maximizing CPU usage across cores. The authors are aware of two main approaches to this: with Dask™ [17] and Spark™ [18, 19]. Dask [17] is a distribution framework capable of distributing generic python workloads. Dask implementations are being built into scanpy [8], though they are still noted on the official scanpy documentation [20] as being “new and highly experimental”, and “many functions in scanpy do not support dask”.

Spark is a popular distributed computing framework that is extremely popular for big data workloads, and has existing tools for ML processing [21]. Pyspark provides a simple user experience on top of Spark allowing for easy integration into python workflows. Though Dask is lighter weight and is very popular for distributing scientific workloads due to its python integration, Spark has very strong backend for many typical mathematical and ML operations. Therefore, while Dask may be a strong contender for distributing existing python code, we think it is worth considering whether rewriting some of these processes with pyspark may benefit from the efficient Spark backend for big data processing.

At the time of writing the authors became aware of another framework for single cell RNA analysis that utilizes Spark, called scSPARKL [22]. Despite the impressive work of Adil et al. [22] to write key functionality of scRNA processing in (py)Spark, we believe our method and findings are worth sharing with the community. In particular, we note that scSPARKL does not offer any support for h5ad files or other common file formats for scRNA data, rather preferring csv in wide format expected to be converted into spark dataframe. The code repository notes that they are no longer actively maintaining the repo and intend to add more file types, but it seems that updates to the repository are quite rare. There are several reasons why we think our work is worth sharing which we list below:

1. We include distributed read from h5ad files for efficient data ingestion, and handle multiple files at once. The h5ad read requires conversion from CSR (Compressed Sparse Row) to COO (COOrdinate) format. Specifically on chunked arrays from CSR format which are not rectangular sub-arrays which adds complexity (See methods).
2. We use a data storage layer (SprayData) that has similarities to the popular AnnData object, but is stored as PySpark DataFrames rather than Pandas Dataframes and SparseArrays. This includes obs, var, as well as new additions for the multi-file nature. These obs and var matrices are not included in scSPARKL [22] to the best of the authors’ knowledge.
3. Implement functions on SprayData objects that are similar to scanpy, but are distributed and scalable, such that a familiar interface for scRNA analysis is maintained for users already familiar with scanpy.
4. We performed analysis over larger datasets and explicitly detail the scaling behavior of cspray over multiple workers, showing that preprocessing is identical to scanpy with the same parameters, and evaluating clustering differences to the commonly used “Leiden” method when using Kmeans (which is more easily distributable).
5. Include a simple LLM based cluster annotation method that can be used to annotate clusters with a pre-defined cell ontology. This is based on the work of Hou et al. [13], with our addition of the cell ontology definition for low resolution clusters and differing prompts, which we describe in the “LLM based cluster annotation” section.

## RESULTS

### cspray: a distributed framework for scRNA processing

cspray is a python codebase, utilizing pyspark (based on Apache Spark [18, 19]) native functionality wherever possible, for processing single cell data in a distributed manner from data ingestion to cell clustering. In order that the functionality is easily transferable to existing python users of the highly popular scanpy [8], we use similar syntax for the main data object holding the data and for subsequent processing steps on it.

The main python object containing the data in cspray is called a SprayData object (Fig. 1a) and like the familiar AnnData [23] object it contains variables X (sparse expression array), var (gene indexing and metadata) and obs (cell indexing and metadata). The key difference in the approach of SprayData is that these variables are stored as PySpark DataFrames rather than Pandas Dataframes and SparseArrays. There are four key indices across these data: the index of a sample, of a gene, of a cell, and of a cluster. We keep indices for cells and genes to only be unique within a sample.

**FIG. 1:**
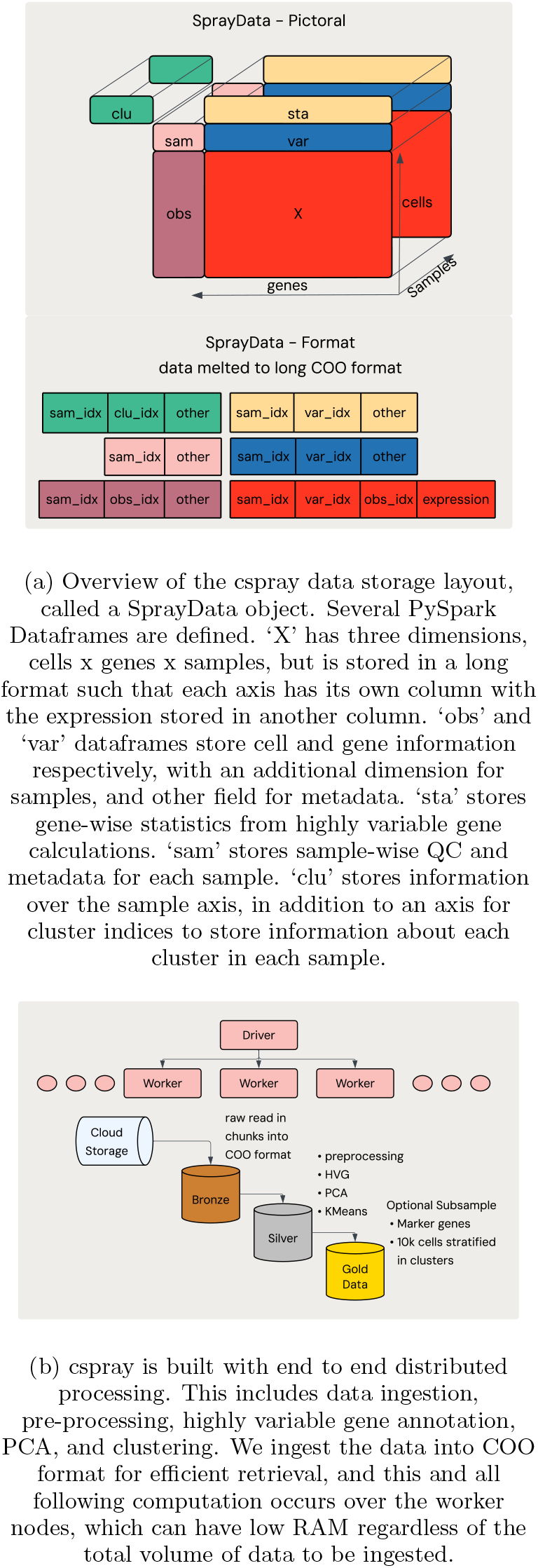
cspray system overview.

Let us address how data is imported and exported from these data structures and the benefits of this approach. We will begin with the ingestion, as this has some complexities as we wish to have the cell by gene data in a long format, which requires some additional logic over a traditional read.

### Data input and output with cspray

Many single cell data are stored in h5ad format, though other formats such as loom and increasingly zarr [24] are used. We have focused on h5ad as we believe it’s the most common, and we can extend from here to other formats. h5ad datasets store the gene expression array in Compressed Sparse Row or Column (CSR, CSC) format, and data is typically stored in chunks on disk. This format stores the elements of the sparse gene expression matrix as an array, together with an array of the index of the column and a smaller array of *pointers* indicating at which entry in the first two arrays data swaps to a new row. This takes less space on disk then Coordinate (COO) format sparse arrays, but COO format data is more easily extensible to multi-sample data. To expand on that, COO data stores three equal sized arrays: one for entries in the matrix, one for column indices, and one for row indices. If one has a table with these three arrays as columns, adding new samples is as simple as appending a new sample to the end of the table, so long as you also have a column indexing the samples.

In our distributed read of h5ad files from disk, we read data from files in chunks, and for each chunk of the CSR matrix that’s read from disk, one must convert it to COO format to place it in a long format table of multiple samples. The challenge here is that the arrays do not correspond to a square submatrix of the original. Rather, they are a contiguous arrays across rows that can span multiple rows. To convert these contiguous row chunks to COO requires one to use the entire pointers array of the matrix to calculate the correct row position in the full sized matrix for each position. For details, see Methods.

We do not currently support reading zarr [24] files, as we focused on what we believe to be the more common h5ad format, but since that format is designed for reading sparse matrices from cloud storage, we expect it to be very efficient and intend to include support for it at a later date.

When one has created the (lazily executed) statement to load h5ad’s from disk, one can save the dataframe to a Delta [25] table. These tables store data in columnar format and apply compression. This compression means that the added data from the conversion from CSR to COO is limited, particularly since Delta uses a range of techniques such as run length encoding, bit-packing and dictionary encoding, which can be effective with repeated values, such as those that occur in indices of sparse arrays.

After ingesting the data into a table on disk, one proceeds with standard scRNA processing.

### Processing at scale and comparison to existing frameworks

Once data is read into a table, we wish to perform actual analysis on it. With cspray, our intention is to build a robust pipeline for processing large amounts of data, in such a way that’s scalable. We expect that cspray would be used to process data to form a basis for users to discover and pick up data at different stages of processing.

Regarding robustness using pyspark, now that the data is ingested, one can follow standard practice for pyspark operations to avoid OOM errors. Most operations, and in particular in-built Spark functionality will handle this automatically, leaving only custom user defined functions (UDFs) and mapping functions as steps with which to be careful. In our case we rely on built-in tooling in pyspark (including mllib) wherever possible, resorting to functionality like ‘mapinarrow’ only where we have to rely on custom logic and where we believe prior grouping strategies mitigate the risk. We also include the option to persist dataframes to disk during execution of functions on the SprayData objects (see Methods). This is not a requirement but can increase overall performance with the downside of increased memory usage on workers. Alternatively, one can save data to disk at intermediate stages during the pipeline. We apply both approaches, especially since we expect data after QC and PCA to be stages that a scientist would likely pick up data for continued analysis.

To test scaling, and comparison of results with standard tooling, we built an end-end pipeline for ingestion followed by the following analysis steps (more details in Methods):

1. filter cells (based on number of genes with expression)
2. filters genes (based on number of cells with expression)
3. filter cells (based on percentage of reads from Mitochondrial genes)
4. (log) normalize expression (log as log(1+p))
5. Highly variable gene detection (Seurat method [7])
6. PCA (mllib [21] implementation per sample)
7. Kmeans (mllib [21] implementation per sample with various k and silhouette score selection)
8. Ranking marker genes (t-test used, though Wilcoxon available)

We ran this pipeline over two sets of data downloaded from the CELL X GENE census [26], a “large” set with twenty samples containing 100k-1M cells, and an imbalanced set with eighteen files with 6k-8k cells and two samples from the large set (details in methods). We then ran our pipeline over these two datasets with 4, 8, or 16 worker nodes, Fig. 2a. The time to perform the end-end processing scales almost linearly with the number of worker machines used, i.e one can scale the speed of ones pipeline simply by adding more workers. This is true for both cases where the data is many large files or where there is more extreme skew (quite common scenarios for sequencing machines). As one would expect, the imbalanced set having fewer cells overall runs faster than the larger set, but more importantly we do not need to wait for larger files to finish as all data is processed at once. Additionally, we note that we observed that all CPU cores are active for most of the compute time.

**FIG. 2:**
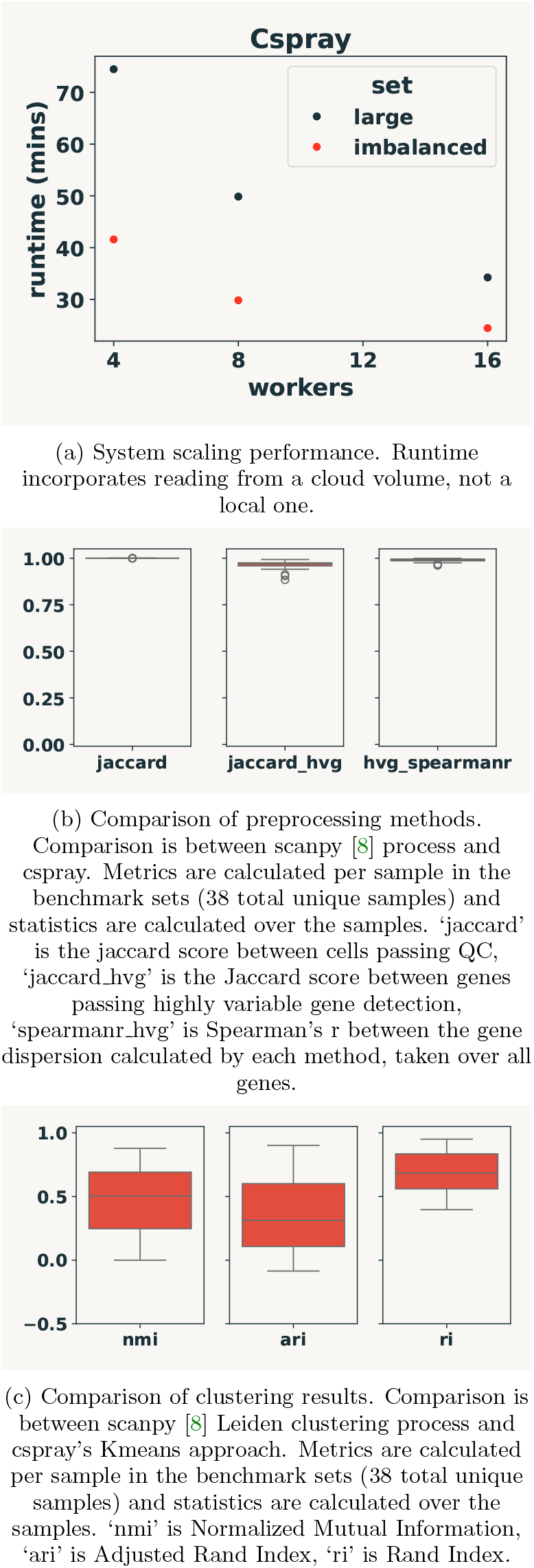
Overview of cspray runtime scaling, and preprocessing and clustering comparisons to industry standard workflow.

Of course, we need to ensure that, other than being robust and scalable, the results match the expected outcome. Since scanpy [8] is a widely used and mature package for single cell analysis we compare the outputs of cspray with it. We include integration tests within the cspray package that run cspray and scanpy on a small sample h5ad file and ensure the ingested array, preprocessing, and variable gene detection match. We also confirmed this over all datasets in the benchmark sets. In particular, we found that preprocessing to filter cells based on counts and mitochondrial reads match exactly, Fig.2b, in terms of Jaccard overlap between the cells kept after filtering. For highly variable gene detection, we must consider that there are small rounding errors as one first performs normalization and logarithm on the expression, then subsequently calculates the dispersion for genes. One can either re-exponentiate or use original values. We chose the latter, but with these multiple repeated calculations, one does expect rounding errors between codebases built on different platforms. The key detail is that these differences should be small, and in terms of ranking genes (the primary goal), should have limited impact. We calculated the spearman correlation over the benchmark sets (Fig.2b) and found that the spearman correlation is close to one for all samples (mean=0.989). In terms of those genes that are placed into the highly variable set, we calculated the Jaccard score between the set of genes that were considered highly variable by scanpy and cspray, Fig.2b, again we see a very strong overlap (mean=0.962), with differences often being genes close to the cutoff.

A commonly used clustering technique in single cell analysis is Leiden clustering [27]. In cspray, we use Kmeans clustering since mllib [21] provides an optimized distributed method for it. We wanted to compare the effect of such a decision, particularly as it relates to clustering at low resolution. We make this resolution decision based on the idea that cspray is designed not for ad-hoc analysis nor identification of niche cell types, but rather for identifying clusters of major cell type groups which should be more robust to a generalized QC pipeline. With that in mind, we calculated three common clustering metrics (Methods) between the cspray Kmeans and the scanpy Leiden [27] approaches. We see that while clustering differs, there is still considerable overlap on the clustering. We note that small changes in the highly variable genes could also have an impact on PCA and clustering.

We note that we included a final optional stage in the cspray package: conversion from the full output into a subsampled set for fast data discovery. Namely, we provide a method to subsample to 10k cells per sample (stratified on the clusters defined in each sample to avoid losing niche clusters), and we also filter only to highly variable genes in each cluster. This creates a much smaller set of data that can be very quickly filtered and visualized for data discovery. Ideally, these data discovery solutions would allow users to search for cell types across these samples, which will be the topic of the next section.

### LLM based cluster annotation

Hou et al. [13] recently demonstrated that LLM based cell type annotation using marker genes at the cluster level was fast and effective, with accuracies of recent LLMs surpassing that of scType [28] and singleR [29]. In brief, the use of LLMs to call cell types in this way proceeds by: passing a prompt with the marker genes of a cluster and the sample tissue to an LLM, which responds with a predicted cell type. Additional strategies can be used to further improve the quality of the cell type calling; Hou et al. [13] used a repeated prompt strategy (majority voting over multiple attempts) and a chain of thought style strategy (using additional biological knowledge in follow up) to improve calling. Hou et al. observed significant variation in performance across tissues, with accuracies from 0.5 - 0.8 with combined scoring of these where a partial match is scored at 0.5 and a full match is 1.0.

Strategies for further improving call type calling via LLMs have been further extended by Ye et al. [14] with their method Large Language Model-based Identifier for Cell Types. Ye et al. [14] explored two other strategies: (i) multi-model integration: a majority voting over multiple attempts, though this uses benchmarking against ground truth to filter the best annotations which would not be feasible when ground truth annotations do not exist, and (ii) talk-to-machine: a more complex strategy that, after initial cell type calling, has continued LLM and data interaction. The “talk-to-machine” strategy is much more applicable in real world systems than their “multi-model integration” as there is no reliance on ground truth, though it does require further aggregations over the large datasets.

The methods of Hou et al. [13] and Ye et al. [14] are a promising direction for scaling cell annotation as cell counts and sample counts continue to rise; however, both require data preprocessing up to and including cell clustering. Our primary interest is to understand if methods like these work on data processed at scale with the same parameters used for preprocessing across all samples rather than one-by-one expert preprocessing and clustering. To that end, we applied a cell type calling framework across our benchmark datasets and compared the annotations to those provided in the original source. In our method for each cluster of cells found by cspray, we pass the 10 marker genes for the cluster to the LLM, together with the tissue of the sample, and format it into our prompt template (see methods). For the LLM, we used GPT 5-1 and employed majority voting over 5 repeated calls. Our prompt contains a list of allowed cell types that can be annotated, and we selected these in such as way as to include cell types that are quite broad, but not exceptionally so. This is performed by searching the cell ontology for cell types that have a range of allowed numbers of descendants and depth within the ontology (for full details see Methods). The reason for this is that our vision for large scale processing and annotation with cspray is that clustering and annotation should both be a high level. That is, not for niche cell types where more generic processing (no expert tuning of parameters per sample) is more likely to fail.

Since we are deliberately limiting the cell type annotations to higher level terms, we do not expect to necessarily get exact matches to existing annotations. We instead focus on the annotations being highly similar or of an ancestor cell type in the ontology. To assess this performance, we use an LLM to compare the labeled annotation of a cluster to the true labels of cells within that cluster. Since the cells in a cluster can have multiple different true annotations, we limit the cell types to compare against to the most common cell types in the cluster (see methods). We manually confirm *post-facto* that the LLM based annotation scoring aligns with our human expectation of if there is a good match. We find that across samples we see an accuracy, on this similarity scoring basis, of 56.3±1.1% across our clusters. Given the breadth of dataset sizes and tissues of origin, we believe this shows that this automated processing and annotation strategy could be used for processing and annotating cells at scale. Further improvements in the specifics of the cell type calling, such as using the “talk-to-machine” strategy of Ye at al., could be employed for additional benefits.

## DISCUSSION

The scale of data in single cell datasets has expanded in some cases to millions of cells per sample, and this should be expected to increase with new technologies. It is therefore important to have methods that can scalably process these data as the scale continues to grow without relying on ever increasing memory on single devices. Beyond standard preprocessing and clustering of data, annotating these data for automated data discovery programs is also key. Namely, being able to identify samples based on the cell types present without requiring expert annotators to have reviewed these datasets.

We introduce cspray, a Spark based package for performing distributed analysis of single cell datasets at scale. The distributed execution of cpray is scalable: simply adding more workers can scale the runtime of the analysis, without requiring any machines that individually have large random access memory. cspray uses a familiar paradigm for processing single cell data, encapsulating the datasets into a single python object with processing operations applied on this object. We support reading the commonly used h5ad file type with distributed ingestion. Data are stored after ingestion in Delta format tables, allowing scientists to collect single or multiple samples of data at various stages of preprocessing to continue ad-hoc analysis if required.

The ability to process very large numbers of large samples poses the question of whether processing many samples with the same parameters, rather than niche ad hoc analysis, leads to useful data output. We suggest that cspray is primarily targeted either at cases where (i) data is so large that traditional methods are unable to cope, or (ii) where one wishes to automatically preprocess new datasets for data discovery with pre-performed standard preprocessing for later niche analysis.

To assess this, we explicitly compare processing with cspray to an industry standard pipeline with scanpy [8]. cspray performs basic pre-processing identically to the scanpy implementation, variable gene detection with almost complete match, and PCA combined with simplistic Kmeans clustering to provide broadly similar clusters to low-resolution clustering with more advanced Leiden [27] clustering routine. Further, this standardized preprocessing and clustering routine with marker gene annotation is sufficient to allow for reasonably accurate reference-free cell type annotation via LLM based cell calling methods.

In summary, cspray is an end-to-end distributed framework for analyzing single cell transcriptomic data that scales well as data volumes grow. The results are very closely matched to industry standard processing tooling for basic workflows, and low resolution clustering and cell annotation perform well enough to enable data discovery across large data sets and ongoing streams from labs.

## METHODS

### Dataset Details

We collected data from the CELL X GENE census (version 2025-01-30). In particular we created two datasets, a set of large files and a set of deliberately imbalanced files. We collected samples with species=homo sapiens, suspension type=cell, is primary data=True. We then selected samples with a raw field in the h5ad object (having both X and var available); for the large set we chose samples with cell count: 100k - 1M, and took every third file to get a set of 20 files. For the imbalanced set, we performed the same filtering except for samples with between 6k and 8k, resulting in 18 files, which we combined with two large files from the large set. The full list of sample ids from CELL X GENE in each dataset are shown in tables I and II.

### Code Details

For full details on the code we suggest referencing our code directly on GitHub. We discuss a few key topics mentioned in the main text below.

#### Sparse Matrix Sub Array Conversion

The cell by gene sparse matrix within an h5ad file is typically stored in CSR format, and the three components of that sparse array (indices, indptr, data) are chunked on disk. Since these can be extremely large, we do not wish to read the entirety of the array in one go to avoid going out-of-memory on the RAM. To avoid this, we can read data in predefined contiguous subarrays from each of these three components. The size of the contiguous arrays can be fixed to ensure good usage of the RAM available on each worker node. In principle, simply reading chunks of data such as these from these files is fairly straightforward.

The main complication is that for handling multiple samples at once, and for later aggregating over the data, the CSR format is less useful than a COO format which nicely matches to a long-format table. The challenge lies in converting a set of contiguous subarrays from the three CSR data components into a set of COO entries for appending into a table of all cell-by-gene entries. This is challenging because, for CSR, contiguous arrays start potentially in the middle of a row of entries in the full matrix and go across the row, looping through successive rows before stopping at some point in a new row. To convert this to COO (absolute column, absolute row, data) one has to calculate what absolute row you started on, and this requires the full indptr array.

To actually implement this we:

- List all files to be read in a Spark dataframe, and for each collect the total number of entries in the matrix
- Create a table with starting and ending indices to read for each file: each file has multiple rows with start and end indices to read contiguous chunks between those indices from disk. The chunk size must be set by the user. This is the one place in cspray where one explicitly sets things to avoid OOM, and one is selecting chunk size for the compute used not for the data. If OOM is hit, it should error early in runtime, preventing wasted compute.
- For ingestion, for each row in the table, we read contiguous chunks (between start and end index) of indices and data, and the entirety of indptr
- We determine the correct row index subarray from indptr based on the start index to match the contiguous subarrays of indices and data
- We construct a CSR array from the contiguous arrays and the extracted indptr subarray
- This sub-CSR array is converted to COO format, and subsequently the row indices are shifted based on the first absolute row value in indptr that corresponds to the contiguous subarrays of indices and data.
- For each row in our table (each is a contiguous chunk range), we apply the above operations using ‘mapinarrow’ which allows the output shape of the dataframe to adjust so that afterwards each row represents one combination of (sample, gene, cell, expression).

#### Dataframe Persistence

There are many functions, representing pre-processing steps, that get applied to the SprayData object that holds the multi-sample expression data. Within these functions, the various elements of the data object (X, obs, var etc) are updated throughout, and there are multiple intermediate dataframes that are created. As one applies these functions, execution with pyspark is lazily executed; i.e. when one writes out one of the dataframes to disk, intermediate dataframes will be used but not cached. This does not pose any issue *per se*, except that if any intermediate dataframes can be reused when one writes the next table out to disk, then caching those intermediate tables can save time during execution. To do this one does need to have sufficient disk (not RAM) space on workers; this is typically much cheaper than adding additional RAM to workers.

We provide a centralized flag in the SprayData object for turning persistence on and off that is then applied within the various functions for preprocessing.

### Benchmark details

We compare with scanpy in its standard usage, very similar to the tutorial code available on their documentation. We do not include steps after marker gene selection (e.g. UMAP) as they are not implemented in cspray as distributed processes. For both scanpy and cspray, to ensure aligned gene name conventions (particularly for mitochondrial gene removal), we join the var (gene) objects to a reference table of the Esnsembl gene ids. This ensures a consistent gene naming convention for each method. For copies of the scanpy and cspray code ran we suggest looking directly at our repo on GitHub.

**TABLE I:** List of sample ids from CELL X GENE in “Large” dataset.

| Sample IDs |
| --- |
| ae4f8ddd-cac9-4172-9681-2175da462f2e |
| 242c6e7f-9016-4048-af70-d631f5eea188 |
| 0ba636a1-4754-4786-a8be-7ab3cf760fd6 |
| 7b55fe5c-d5c8-48e0-a1a3-54c5b9074f3f |
| 80a2c5b6-02e7-4fc0-9f12-179f5247c1bc |
| 43245158-5ae1-4e71-a9a6-67eef49c26bc |
| b252b015-b488-4d5c-b16e-968c13e48a2c |
| 21d3e683-80a4-4d9b-bc89-ebb2df513dde |
| 350237e0-9f48-4cbd-9140-3b44495549f3 |
| ed5d841d-6346-47d4-ab2f-7119ad7e3a35 |
| 1b9d8702-5af8-4142-85ed-020eb06ec4f6 |
| 576f193c-75d0-4a11-bd25-8676587e6dc2 |
| 55003f67-c494-46f1-83fb-902745646379 |
| 18e2a8c5-33f7-455e-a58a-b2ba6921db27 |
| fd072bc3-2dfb-46f8-b4e3-467cb3223182 |
| 7f7faf6b-f11d-4f07-bc1c-188a4472748d |
| 518d9049-2a76-44f8-8abc-1e2b59ab5ba1 |
| 9fcb0b73-c734-40a5-be9c-ace7eea401c9 |
| cd2f23c1-ae1f-48ae-8eb4-0bcf124e567d |
| 76150f40-1989-4977-9e23-696e72d59d9e |

#### Benchmarking: preprocessing stages and parameters

The preprocessing stages and parameters used are listed below (note that where not mentioned the default parameters of scanpy are used as per v1.11.4)

- **Data Ingestion**:
  - h5ad files from CELLxGENE
- **Cell Filtering**:
  - min genes = 100
- **Gene Filtering**:
  - min cells = 3
- **Cell Filtering: Mitochondrial Reads**:
  - gene name prefix = ‘MT-’
  - remove cells with *>*8% mitochondrial reads
- **Normalization**:
  - 10,000 counts per cell
- **Log Transformation**:
  - log1p (*y* = log(1 + *x*))
- **Highly Variable Gene Selection**:
  - method: flavor = ‘seurat’ (per [7])
  - n top genes = 1000

**TABLE II:**
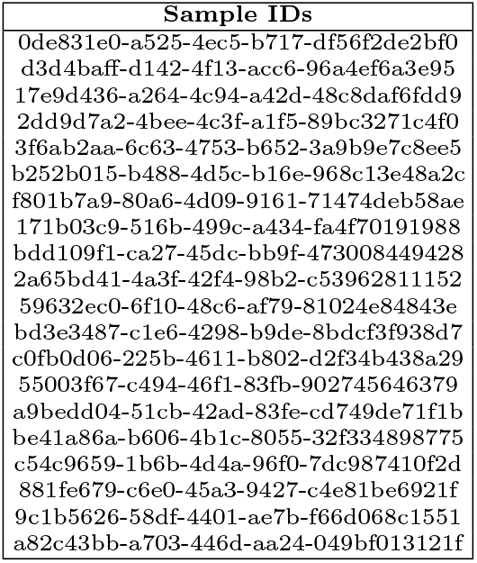
List of sample ids from CELL X GENE in “Imbalanced” dataset.

#### Benchmarking: PCA, clustering, marker genes

We use pyspark MLlib [21] PCA and Kmeans implementations for dimension reduction and clustering. These are already extremely well distributed making them an excellent choice. Swapping to other MLlib clustering methods would be fairly simple, and could be a future extension. We also tried distributing scanpy methods for PCA and clustering, on the basis that after pre-processing and HVG selection, the total data volume per sample is massively reduced. While this works, we already started running into issues for large files even after the data compression from PCA and HVG, and so we chose to use pyspark native approaches throughout as the default.

PCA and clustering with MLlib operate on a per file basis and are distributed for that execution, but they run samples in serial. This is due to the implementation with MLlib and the inability to use these methods following a groupby operation. Nevertheless, the implementation retains good usage of the cores across the system during these operations. PCA is expected to perform very similarly to the scanpy implementation since they rely on well tested PCA implementations from major libraries, albeit with differing random seeds. For our KMeans clustering approach we try 2,3,4, or 5 clusters, and calculate the Silhouette score for each. The best performing clustering ‘k’ labels are placed into a final ‘cluster’ column as the final labels for clusters.

Once clusters are labeled, we perform marker gene calculation (10 per cluster are stored by default and in benchmarking). The (log) normalized expression is compared for each gene across cells in a cluster vs. all others within the sample. We provide both ‘t-test’ and ‘Wilcoxon’ (Mann-Whitney U) methods, and we used the t-test method in the scaling analysis of cspray.

Parameter choices for PCA and clustering are below:

- **PCA**:
  - n comps = 50
- **Clustering**:
  - cspray: kmeans (2,3,4,5); best selected by silhouette score
  - scanpy: Leiden at resolution=0.05, with n iterations=2, flavor=igraph

#### Benchmarking: metrics

After doing the processing and clustering, comparison with scanpy was performed. To do this the datasets for each sample were joined on either the cell barcode, or the gene index (ensembl id). The Jaccard index was calculated between the sets of cell barcodes in the output from scanpy and those from cspray, in all cases a value of 1.0 was obtained. For highly variable genes, the Jaccard index was calculated between the sets of genes in the detected 1000 highly variable genes, and additionally the spearman correlation coefficient was calculated (using scipy [30] stats.spearmanr method) over all genes across the sample (we note that an inner join was performed).

We compare the cluster labeling of scanpy at low leiden resolution with cspray, using kmeans with 2,3,4, or 5 clusters (best selected by silhouette score). For clustering comparison we compare to scanpy with Leiden clustering since that method is commonly used in the examples and real world implementations of scanpy. We use Kmeans clustering, and therefore would not expect an exact match to the scanpy output. Nevertheless, we ask what level of similarity there is between Kmeans clustering with low numbers of clusters and Leiden clustering at low resolution. Three metrics are calculated: normalized mutual information (NMI), Rand Index (RI), and Adjusted Rand Index (ARI). In particular the output from each method are joined on cell barcode, for each sample, and then the three metrics are performed over the cluster columns, using the scoring functions for each method available in scikit-learn [31].

For cspray we only include options for 2 or more clusters since a lot of the following analysis would currently fail with a single cluster. We note that for two samples (0de831e0-a525-4ec5-b717-df56f2de2bf0 and c0fb0d06-225b-4611-b802-d2f34b438a29) scanpy only gave a single cluster. In this case NMI and ARI are identically 0 by definition, though RI took values 0.42 and 0.89.

#### Benchmarking: Compute used

We used machines (driver and workers) with 32GiB RAM, 8 cores, and temporary storage of 300GiB. Dependencies are as detailed in the package’s install script, pyproject.toml, and we note that for benchmarking the scripts were run on Databricks Runtime 16.4 for cspray and 15.4 for scanpy, with scanpy version 1.11.4. For cspray, we additionally allowed an autoscaling local storage on worker nodes, that is used if cached/persisted data grows above the temporary storage of the workers. This can sometimes be advantageous and cheaper than using larger compute. Data persistence is optional with cspray though benchmarks were run with it enabled; local storage scaled for the large benchmark set but not for the imbalanced set. We note that some files required *>*200GiB RAM machines to process those with scanpy. The Spark engine used was ‘Photon’-enabled, though the same code runs with the open source Spark engine, albeit slower by a factor of 1.998.

#### Cluster cell type calling

We began by defining the cell types from the OBO foundry cell ontology [32] that were neither so high level as to be non informative, and not so niche that low resolution clustering marker genes would be unlikely to call accurately. Specifically we select nodes with fewer then 60 total descendants (too niche), or is over 5 edges from the eukaryotic cell node (too deep).

We used GPT-5-1 as the LLM for cell type calling, with majority voting over 5 repeated calls. The full prompt used is included below, and it was passed the tissue (determined as the modal tissue across all cells in the sample) and the marker genes (10 per cluster) for each cluster.

Prompt:

You will be given a list of genes that are most differentially expressed in a cluster of cells in a scRNA analysis as well as a tissue type the cells are from. Your task is to give the cell type label from the list below. You MUST only provide your final answer and no further information. e.g “glial cell” or “fibroblast” NEVER any thoughts or discussion

Assign a cell type from one of these: [‘T cell’, ‘endothelial cell of vascular tree’, ‘retinal cell’, ‘visual system neuron’, ‘ecto-epithelial cell’, ‘GABAergic interneuron’, ‘neurecto-epithelial cell’, ‘stem cell’, ‘macrophage’, ‘epithelial cell of nephron’, ‘sensory neuron’, ‘connective tissue cell’, ‘mononuclear phagocyte’, ‘kidney medulla cell’, ‘smooth muscle cell’, ‘neuron associated cell’, ‘stromal cell’, ‘columnar/cuboidal epithelial cell’, ‘blood vessel endothelial cell’, ‘mature B cell’, ‘endo-epithelial cell’, ‘professional antigen presenting cell’, ‘cerebral cortex neuron’, ‘meso-epithelial cell’, ‘neuron of the forebrain’, ‘fibroblast’, ‘myeloid cell’, ‘kidney epithelial cell’, ‘afferent neuron’, ‘kidney cell’, None, ‘striated muscle cell’, ‘epithelial cell of alimentary canal’, ‘retinal ganglion cell’, ‘lymphocyte of B lineage’, ‘interneuron’, ‘macroglial cell’, ‘intestinal epithelial cell’, ‘muscle cell’, ‘hematopoietic lineage restricted progenitor cell’, ‘myeloid leukocyte’, ‘inhibitory interneuron’, ‘glial cell’, ‘glutamatergic neuron’, ‘visceral muscle cell’, ‘phagocyte (sensu Vertebrata)’, ‘B cell’, ‘endothelial cell’, ‘dendritic cell’, ‘alpha-beta T cell’, ‘non-striated muscle cell’, ‘glandular secretory epithelial cell’, ‘endocrine cell’, ‘GABAergic neuron’, ‘progenitor cell’, ‘kidney cortical cell’, ‘conventional dendritic cell’, ‘peripheral nervous system neuron’, ‘secretory epithelial cell’, ‘autonomic neuron’, ‘efferent neuron’, ‘cardiocyte’, ‘sensory receptor cell’, ‘lymphocyte of B lineage, CD19-positive’, ‘mature T cell’]

To this system prompt, for each cluster the following message is sent as the user “Tissue= {tissue}, list of marker genes = {marker genes} “.

In order to assess the cell type calling performance, we first note that we must consider that we label cells using a subset of the ontology. In particular, we wish for cell types to either be the same, further up the ontology tree, or broadly accepted as being a very similar cell type. Additionally, for each cell in a cspray defined cluster, therefore with the same cell label, they can each have different labels from the reference set. To adjust for this, we identified a list of reference cell types in each cspray defined cluster, in particular we keep the largest group, second largest group, etc until more than 66% of cells in the cluster have a reference definition. We then compare the cspray cell label with those labels from the reference set. Our primary comparison is via an LLM-as-a-judge [33] system to assess if the cluster cell label, and those from the reference set are equivalent in our setup, the prompt used was:

You are an expert in single cell labeling. A user has clustered some scRNA data and performed cell labeling, and in particular they have taken a cell ontology and removed,more niche cell types in order to focus on the major cell types (ie they cut the tree). They additionally have a set of “true” labels provided by someone else, however within each of the users clusters there can be several cell types as different by the “true” labels. When there are more than one, that means that the user ordered them by prevalence and included them until they reached >66 percent of cells in the cluster represented in the comma separated list.

The users wishes to evaluate whether their label is equivalent, ie the same cell type or a cell type higher in the tree, to the true label. If the users label matches (in the sense suggested in the last sentence) the any of the true labels, then you should return “True”, otherwise return “False”. If the true cell type is ‘unknown’ you can output “True”.

We further validated that those judgements aligned with our expectations (*post-facto*). We also note that this analysis was performed over all unique samples across the two benchmark sets. In the final reported performance of 56.3±1.1%, the error term is the standard deviation over 5 repeated evaluations.

## CODE AND DATA AVAILABILITY

Cspray is available on GitHub at https://github.com/databricks-solutions/cspray. The data used for the benchmarking is from cell x gene census, available under a CC BY 4.0 license, and we provide a full list of the datasets used in the manuscript in the methods section.

## ACKNOWLEDGMENTS

The authors are grateful for the support of Ramachandran Venkat, Jon Forinash, and Alex Barreto.

## AUTHOR CONTRIBUTIONS

PGH conceptualized the study. PGH wrote the cspray code with input and discussion from MF and EMS. All authors designed the benchmarks. PGH wrote and ran the benchmarks, and prepared the figures. All authors interpreted the results. PGH wrote the manuscript with input from all authors.

